# Co-editing *PINK1* and *DJ-1* genes via AAV-delivered CRISPR/Cas9 system in adult monkey brains elicits classic Parkinsonian phenotypes

**DOI:** 10.1101/2020.09.19.305003

**Authors:** Hao Li, Shihao Wu, Xia Ma, Xiao Li, Tianlin Cheng, Zhifang Chen, Jing Wu, Longbao Lv, Ling Li, Liqi Xu, Wenchao Wang, Yingzhou Hu, Haisong Jiang, Yong Yin, Zilong Qiu, Xintian Hu

## Abstract

Whether direct manipulation of Parkinson’s disease (PD) risk genes in monkey brain can elicit Parkinsonian phenotypes remains an unsolved issue. Here, we employed an adeno-associated virus (AAV)-delivered CRISPR/Cas9 system to directly co-edit *PINK1* and *DJ-1* genes in the substantia nigra (SN) region of four adult monkey brains. After the operation, two of the monkeys exhibited all classic PD symptoms, including bradykinesia, tremor, and postural instability, accompanied by severe nigral dopaminergic neuron loss (over 60%) and α-synuclein pathology. The aged monkeys were more vulnerable to gene editing by showing faster PD progression, higher final total PD scores, and severer pathologic changes compared with their younger counterparts, suggesting both the genetic and aging factors played important roles in PD development. This gene editing system can be used to develop a large quantity of genetically edited PD monkeys over a short period, thus providing a practical transgenic monkey model for future PD studies.

## Introduction

Parkinson’s disease (PD) with a prevalence of 2-3% in humans above 65 years old is the second-most common neurodegenerative disorder (1). It is typically characterized by a cluster of specific motor symptoms, including bradykinesia, rigidity, postural instability and tremor, and by a set of pathologic hallmarks, including severe dopaminergic neuronal loss in the substantia nigra pars compacta (SNpc) and intracellular inclusions containing aggregates of α-synuclein, which are termed as Lewy bodies (LBs) or Lewy neurites (LNs) (1, 2). The severe dopaminergic neuronal loss which appears in every PD patient is the most important hallmark, and the Lewy body and Lewy neurite appear in 77% to 95% patients according to different studies (3, 4). Unfortunately, limited by its unclear etiology, there is no effective treatment to alleviate PD progression and no preventative measure currently (5), and thus, there is a significant increase of PD patients every year (~0.02%) (6). To solve this ongoing PD crisis, early diagnose and prevention of this disease has become a consensus in the field of PD treatment. Etiological PD models, such as transgenic models, which can mimic the development of progressive process of PD are the foundation to achieve this goal. Unfortunately, there is no breakthrough being reported in this area yet.

In addition to etiological PD models, selecting proper animal to carry out the modeling is equally important. Non-human primates are ideal experimental animals to study the etiology and pathogenesis of PD due to their evolutionary proximity, physiological similarity, close aging process, and the equivalent behavioral symptoms and pathological changes as PD patients (7, 8). More importantly, our recent study showed that they can develop PD naturally. Unlike the existing traditional monkey PD models, which are induced by neurotoxins, including 1-methyl-4-phenyl-1,2,3,6-tetrahydropyridine (MPTP) or 6-hydroxydopamine (6-OHDA) models, only mimic the symptoms and partial pathology hallmarks of PD, the gene-edited PD monkey model, which is established by changing monkey genes to mimic the PD genes dysfunction discovered in human patients, is an etiological model. It can be used to study the pathogenesis and screen the early biomarkers of PD (9), and will provide new insights into PD early diagnose, prevention and future gene therapy. To date, however, no successful transgenic PD monkeys have been reported, despite decades of research recapitulating the genetic factors involved in PD etiology and pathogenesis (10, 11). Nevertheless, recent progress in gene editing technology is heralding a new era in the development of genetically manipulated PD monkeys (12–16). Especially the clustered regularly interspaced short palindromic repeat/CRISPR-associated protein 9 (CRISPR/Cas9) system allows sequence-specific gene editing in many organisms and provides a valuable tool to generate gene-editing models of human diseases (17–20).

It is well known that PD risk gene mutations are the primary causes of neurodegeneration in early onset PD patients (11). Notably, *PINK1/PARK6* gene is an autosomal recessive PD risk gene implicated in mitochondrial functions, and the *PINK1* mutations account for 1%–7% of early-onset PD (21). One critical function of the PINK1 protein is the phosphorylation of Parkin protein (an E3 ubiquitin ligase), which is important for protein degradation in proteasomes (10, 21). However, the loss-of-function mutations of *PINK1 in vivo* cannot elicit PD clinical symptoms in monkeys (16), and our data revealed that only editing *PINK1* gene via adeno-associated virus (AAV)-delivered CRISPR/Cas9 system on monkeys’ SN region could not elicit PD clinical symptoms either for it only caused about 40% nigral dopaminergic cell loss [to elicit PD symptoms, the dopaminergic cell loss have to be above 50% (22)] (**Supplementary Fig. S1A**). Therefore, just *PINK1* dysfunction is not enough to trigger PD development in monkeys. Meanwhile, another autosomal recessive PD risk gene, *DJ-1/PARK7*, which encodes an antioxidant protein, potentially is associated with *PINK1* mentioned above to inhibit oxidative stress and α-synuclein aggregations in PD (23, 24). The second experiment was carried out in our lab in which the *DJ-1* gene in the SN region of the monkey brain was also edited via the AAV-delivered CRISPR/Cas9 system. The data showed that this operation could not elicit PD clinical symptoms either for it also caused only about 40% nigral dopaminergic cell loss (**Supplementary Fig. S1B**).

In this study, based on the results from the two sets of preliminary data, we used the AAV-delivered CRISPR/Cas9 system to co-edit the *PINK1* and *DJ-1* genes in monkey brains to investigate the possibility that classic PD symptoms might be elicited by this double editing strategy.

## Results

### Virus induced mutation efficiency tests in vitro

We first designed two small-guided RNAs (sgRNAs) targeting rhesus monkey *PINK1/PARK6* (sgRNA-PINK1-A, sgRNA-PINK1-B) and *DJ-1/PARK7* (sgRNA-DJ-1-A, sgRNA-DJ-1-B) (**Fig. 1A, B, C, D**) respectively, and then tested the efficiency of the sgRNAs in COS7 cell lines from African green monkeys. After transfection of the SaCas9 and sgRNA vectors into COS7, we collected the cells and examined whether CRISPR/Cas9 induced genetic mutations in the sgRNA-targeting regions by polymerase chain reaction (PCR) amplification. We found that all the sgRNA-PINK1-A, sgRNA-PINK1-B, sgRNA-DJ-1-A, and sgRNA-DJ-1-B had effectively caused missense mutations in the protein coding regions of the COS7 cells (**Supplementary Fig. S2**). To co-edit both the *PINK1/PARK6* and *DJ-1/PARK7*, we co-injected four AAVs (AAV9-Syn-SaCas9-sgRNA-PINK1-A/B and AAV9-Syn-SaCas9-sgRNA-DJ-1-A/B) into one side of the SNs of the monkey’s brain, along with an AAV-Seen-GFP to label the injection sites.

**Figure 1.**
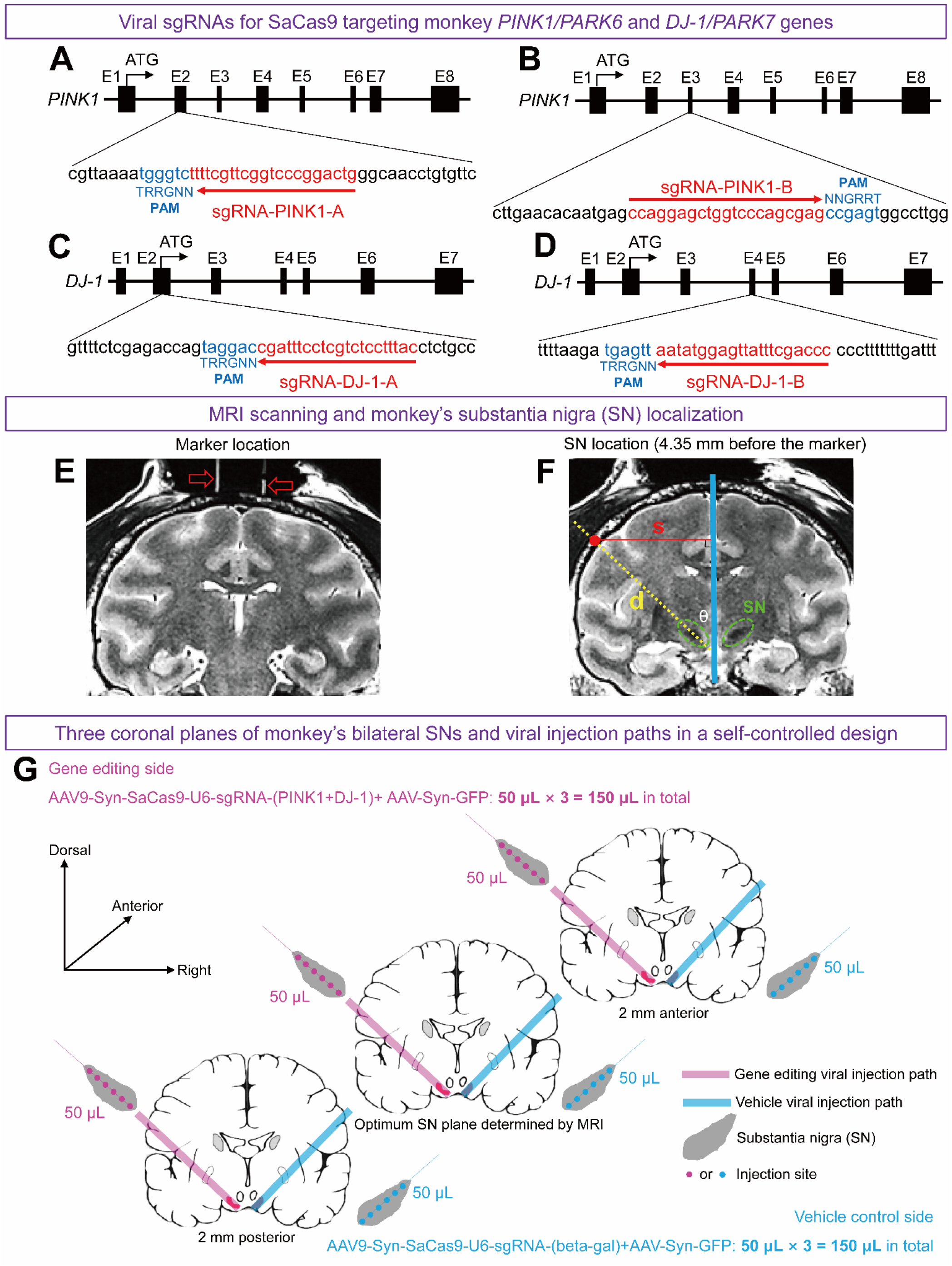
Viral design of sgRNAs for SaCas9 gene editing systems and magnetic resonance imaging (MRI) localization for viral injection paths of the substantia nigra (SN) in monkey’s brain. (**A-D**) SaCas9-corresponding sgRNAs were designed for monkey *PINK1/PARK6* and *DJ-1/PARK7* genes. The targeting sites of *PINK1* and *DJ-1* in rhesus monkey genome are shown in A-B and C-D respectively, in which the sgRNA binding sequences are highlighted in red and PAM sequences are marked in blue. (**E**) The marker (the two glass tubes filled with glycerin indicated by red hallow arrows) implanted in the monkey Old1’s skull was used as the reference to localize the SN in the coronal section. (**F**) The location of the monkey Old1’s SN (green circles) and the injection path of one side of the monkey Old1’s brain (dashed yellow line) are illustrated. The blue line is the symmetric axis of the monkey brain’s coronal section. The red dot indicates the location of the injection hole in the skull. The **s** is the horizontal distance from red dot to the blue line and the **d** is the length from the skull to the SN. The **θ** is the angle between the yellow line and blue line (°), which can be calculated by ****sin θ = s / d****. With both the **s** and **θ** values, the red dot on the skull will be accurately localized. (**G**) Three parallel paths for viral injections containing 18 injection sites which covered the left SN are illustrated in pink lines, and three parallel paths of AAV-vehicle control injections containing 18 injection sites which covers the right SN region are illustrated in blue lines.

### Parkinsonian symptoms assessments

In this study, to minimize the influence of individual difference, a self-controlled design to develop hemi-Parkinsonism monkey model was used in which four male adult rhesus monkeys were involved. As age is an important risk factor in PD pathogenesis (25), the four monkeys were divided into two age groups: an old monkey group (Old 1 and Old 2: 21.5 ± 2.12 years old) and a young monkey group (Young 1 and Young 2: 10 ± 0 years old).

In hemi-Parkinsonism monkey model, the behavioral phenotypes are sensitively to be measured, including the apomorphine-induced rotation and hand preference test. Here, to ensure a whole coverage of viral injections in the targeted SN, we performed the AAV injections into the targeted side of the SNs of each monkey brain through three oblique parallel pathways, which had been determined by a unique magnetic resonance imaging (MRI)-guided high accuracy localization procedure (error < 1 mm) previously developed in our lab (26, 27) (**Fig. 1E-G** & **Supplementary Table S2**). Six weeks after the viral injections, the total PD score of each monkey was marked by the improved version of *Kurlan* scale (*a Monkey-Parkinsonism Rating Scale*) (28), which has been widely used in PD monkey studies (29). The average total PD scores of all the gene edited monkeys progressively increased over time (**Fig. 2A**). The increase of total PD scores over time of each individual monkey was illustrated in **Fig. 2B** and those of the two old monkeys were significantly higher. We found that the old monkeys eventually developed to typical PD with the total PD scores were 14 (Old 1) and 8 (Old 2), and the disease progression was significantly faster and severer than the young group. On the other hand, the young monkeys just developed to mild PD with the total PD scores were 5 (Young 1) and 6 (Young 2), and the disease progression was much slower and slighter accordingly.

**Figure 2.**
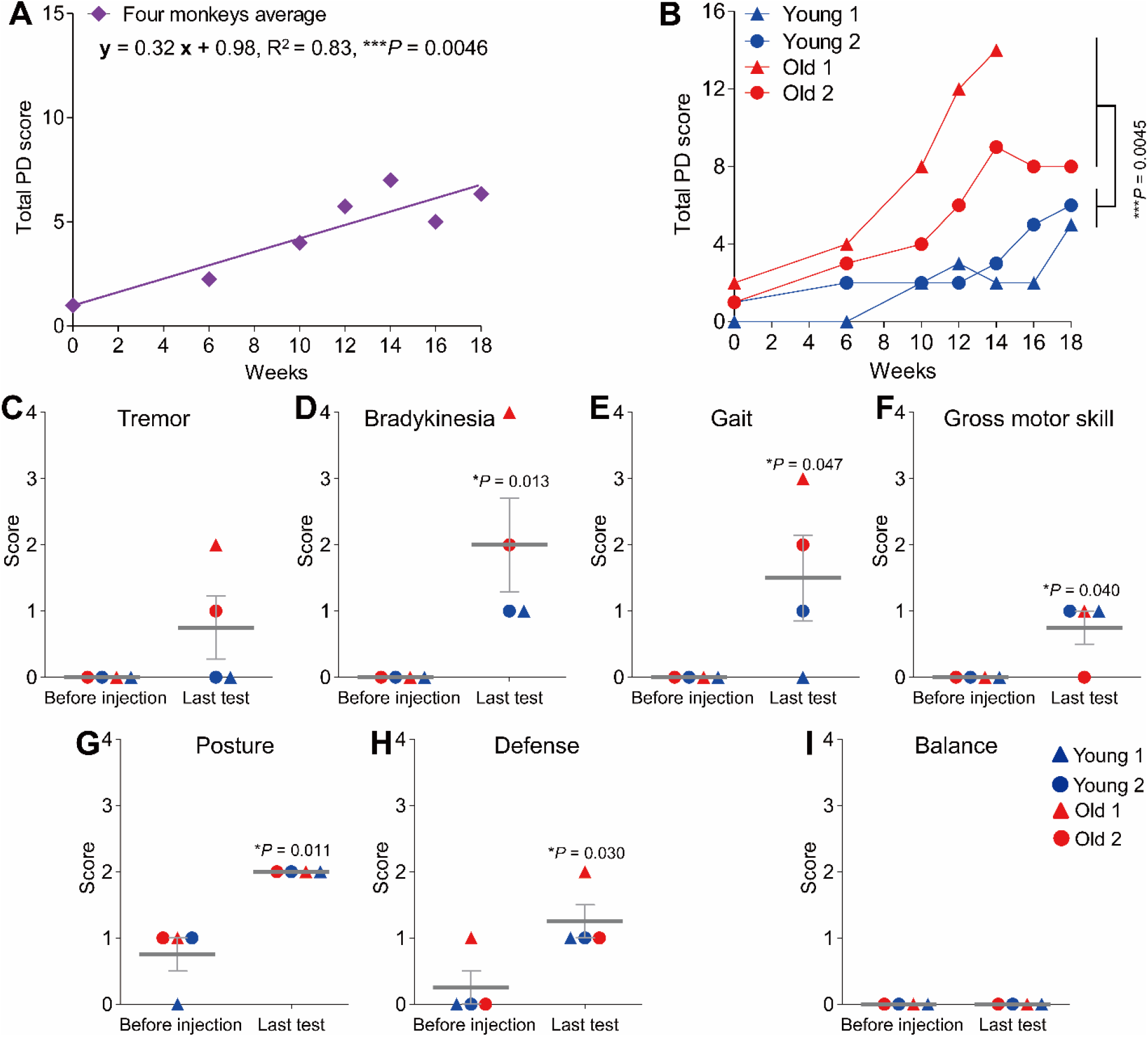
Assessments of diagnostic Parkinsonian symptoms induced by AAV-delivered CRISPR/Cas9 editing of *PINK1* and *DJ-1* genes in monkey SN using recorded daily behavior. (**A**) Averaged total PD score of the four adult monkeys quantified by the improved *Kurlan* scale progressively increased after *PINK1* & *DJ-1* genes editing. *P* value means a very significant correlation of the total PD score and time (weeks), becomingly mimicking the characteristics of PD progression. (**B**) The total PD score of each monkey was also increased with time (weeks), and the scores of the old monkeys were always higher and significantly severer than that of the young monkeys (monkey Old1 died after the data collection of 14^th^ week). (**C-I**) Changes of score for each item consisted the improved *Kurlan* scale between the baseline (before the viral injection) and the last test (the data of monkey Old 1 was from the14th week, whereas the others were from the 18th week). (**C-F**) Tremor, Bradykinesia, Gait, and Gross Motor Skill, which are the core symptoms in PD diagnosis amount to PD monkeys, all increased from a baseline of zero. (**G-H**) Posture and Defense increased significantly compared with the baseline. (**J**) Balance, which usually appears at late stage of PD monkeys, did not change. Data in (C-I) are means ± SEM.

The next are the changes of the scores of the seven items that make up the part A of the improved *Kurlan* scale at the beginning and the end of the experiment. The seven PD-related behavioral items, including “Tremor”, “Bradykinesia”, “Gait”, “Defense”, “Posture”, “Balance” and “Gross motor skills”, which were not observed or very low at the commencement of the experiment, all appeared or increased at the last test except for the “Balance”, which usually appears at late stage in PD monkeys (**Fig. 2C-I**). According to the UK Parkinson’s Disease Society Brain Bank clinical diagnostic criteria (11), core clinical symptoms in PD patients include the essential criteria “bradykinesia”, and at least one of the following: resting tremor, postural instability (which is reflected as the gait, and gross motor skill disabilities in the monkey’s scale, **Supplementary Table S3**). Therefore, the above data almost perfectly matched the diagnostic criteria of PD in humans (**Supplementary Movie S1, S2**). That is, those gene edited monkeys were Parkinsonism. Again, like the total PD score (the sum of the scores of the seven individual items), the scores of the individual items in the old group were generally higher than those of the young group (**Fig. 2C, D, E**), indicating their greater sensitivity to the gene editing operation.

Following the classic PD symptoms assessments, a pharmacological validation was carried out. Compared with the symptom validation, which is a relative rough quantification of the animal’s daily behavior, a pharmacological validation is a better controlled and more PD specific quantitative test. Here, we measured PD-related dopaminergic system dysfunction using a classic pharmacological rotation test paradigm, in which the monkey is administrated with apomorphine (Apo), an agonist of dopamine D1 receptor. As the postsynaptic dopaminergic receptors in the striatum of lesioned side are always more sensitive to Apo than those in the healthy side are, this imbalance would cause the monkey to rotate to the healthy side after the Apo administration (30, 31). After the viral injections and prior to the administration of Apo, the monkeys slowly developed a tendency to rotate to the lesioned side because of the damage caused by the gene editing (**Fig. 3A**). On the other hand, the rotation was reversed to the healthy side after the administration of Apo (**Fig. 3B**). Both data indicated a significant impairment of the dopaminergic system in the monkey brains by the gene editing (**Supplementary Movie S3**). Interestingly, rotations were predominantly identified in the young monkeys both in the spontaneous and Apo-induced conditions, which were significantly more than those in the old monkeys (**Fig. 3C-D**), indicting the young individuals were more sensitive to the rotational tests which has been previously demonstrated in different aged animal groups (32, 33).

**Figure 3.**
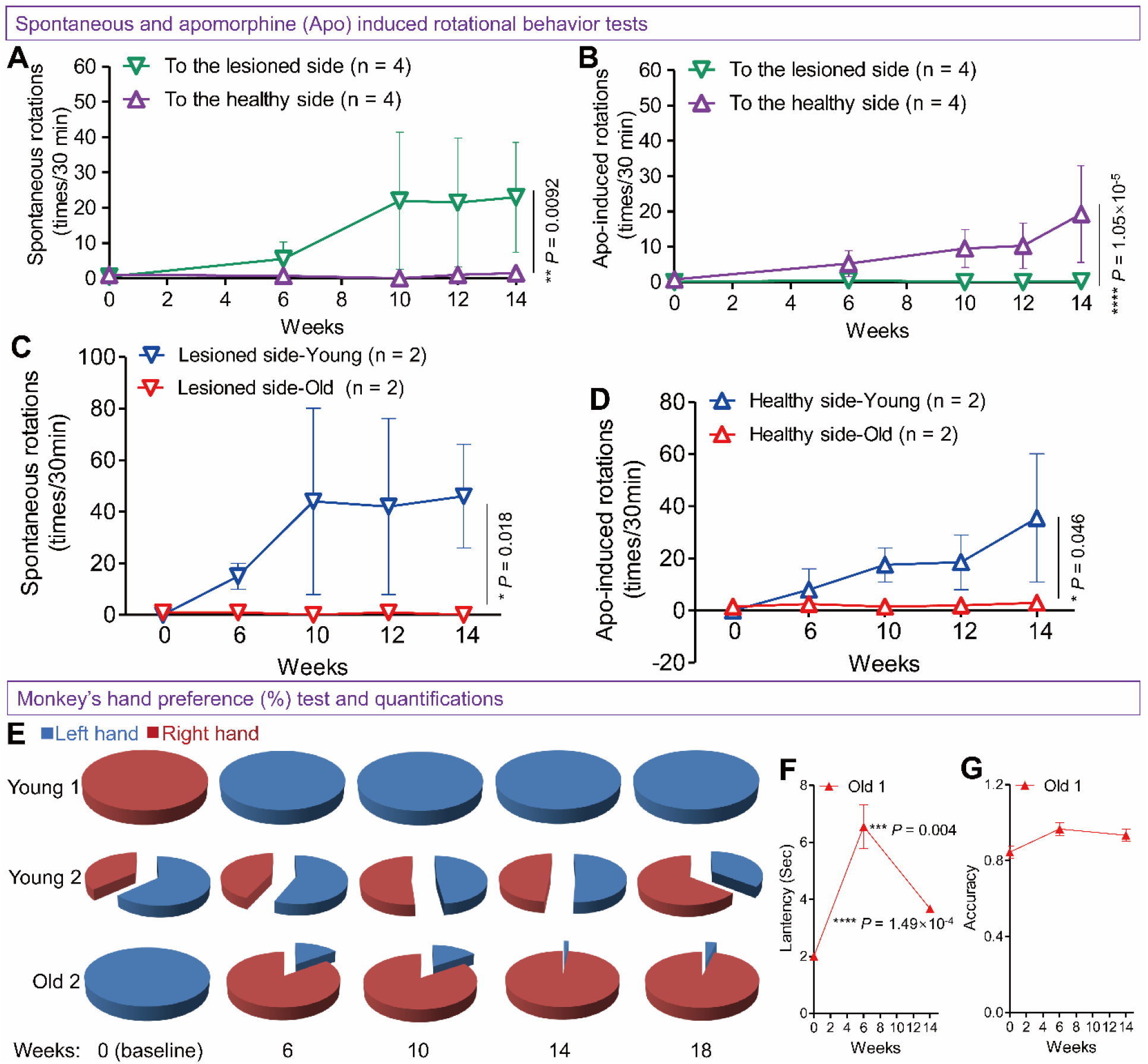
Quantitative assessments of the nigral dopaminergic system functional deficits of *PINK1* and *DJ-1* edited monkeys using spontaneous and apomorphine (Apo)-induced rotational behavior tests and the fine motor skill task. (**A**) Spontaneous rotational behavior was observed in a progressive manner after the gene editing operations and the quantifications indicated that the monkeys were significantly rotate to the lesioned side. (**B**) Apomorphine (Apo)-induced rotational behavior was also observed in a progressive manner after the gene-editing operations and the quantifications indicated that the monkeys were significantly rotate reversely to the healthy side. (**C**) Comparison of the spontaneous rotational behavior between the young and old groups. The young group showed significantly more spontaneous rotations to the lesioned side than the old group. (**D**) Comparison of the Apo-induced rotational behavior between the young and old groups. The young group again showed significantly more Apo-induced rotations to the healthy side than the old group. (**E**) Three monkeys obviously changed their hand preference after the gene editing, but the monkey Old 1 did not. (**F**) monkey Old 1 displayed a significant increase in the latency during fine motor skill task. (**G**) Accuracy of monkey Old 1 during fine motor skill task did not change compared with the baseline. Data in (A-D) and (F-G) are means ± SEM.

Fine movement dysfunction is another common behavioral deficit in PD patients and can be tested in non-human primates using a food grasping task. Before this test, the lesioned side of SNs was selected based on the hand preference of each monkey: Young 1 and Old 1 were 100% right-handed, so AAVs-Cas9 targeting the *PINK1* and *DJ-1* genes was injected into their left SNs. Conversely, monkey Young 2 and Old 2 were mostly left-handed, and thus the genes in their right SNs were edited. After the viral injections, three out of the four monkeys switched their preferred hand in the food grasping task (**Fig. 3E**), clearly demonstrating a fine movement deficit in their preferred hand because they were unable to use their previously preferred hand to grasp food effectively and had to switch to the other (**Supplementary Movie S4**). The latencies of the three monkeys who changed their hand preference were significantly increased (**Supplementary Fig. S3A**), accompanied by decreased accuracy (**Supplementary Fig. S3B**). Only the monkey Old 1 maintained its hand preference, but with a significantly increased latency (**Fig. 3F**) and unchanged accuracy (**Fig. 3G**), revealing a less severe form of the fine movement deficits.

In summary, the strategy of co-editing *PINK1* & *DJ-1* genes by AAV-delivered CRISPR/Cas9 systems in unilateral SNs of adult monkeys can progressively elicit typical Parkinsonian symptoms.

### Pathological hallmarks assessments

Three monkeys were sacrificed for PD pathologic tests. The brains of monkey Young 1 (10 years old) and Old 1 (23 years old) were used for the classic PD pathology examination, whereas the brain of monkey Young 2 (10 years old) was used for viral transfection and gene-editing consequence tests.

The key pathology hallmark of PD is the severe loss of dopaminergic neurons and obvious morphological changes of the survived dopaminergic neurons in the substantia nigra pars compacta (SNpc) (4, 11). Tyrosine hydroxylase (TH) immunostaining, which is a classic staining method for visualization dopaminergic neurons, showed that the genetically edited SN side became pale and blurred under lower resolution images (**Fig. 4A-B**), and the cellular morphologies of the nigral TH positive neurons (dopaminergic cells) were severely deformed compared with the those of the AAV-vehicle control side under higher resolution images (**Fig. 4C-D**). Further detailed observations indicated that the somata of the TH positive neurons in the genetically edited SN side were not clearly visible (**Fig. 4E-F,** solid blue arrow), and the neurites were almost lost completely (**Fig. 4E-F,** hollow blue arrow). Cell count and statistical analysis revealed that over 60% of the TH positive cells (average of the two monkeys) were lost in the genetically edited side compared with those in the AAV-vehicle control side (**Fig. 4G**). Further assessments indicated that the TH positive cells were all significantly lost over 50 % in the genetically edited SN side compared with those in the AAV-vehicle control side both in monkey Old1 (**Fig. 4H,** about 64% loss) and Young 1 (**Fig. 4I,** about 58% loss). Importantly, the TH positive cells on the AAV-vehicle control side were not significantly different from those of normal controls either in the morphology or cell count (**Supplementary Fig. S4**), indicating that the AAV-vehicle control did not cause any observable lesions to the SN. Therefore, the CRISPR/Cas9-edited monkeys successfully mimicked the nigral dopaminergic cell loss of PD, which is the most important pathologic hallmarks of PD.

**Figure 4.**
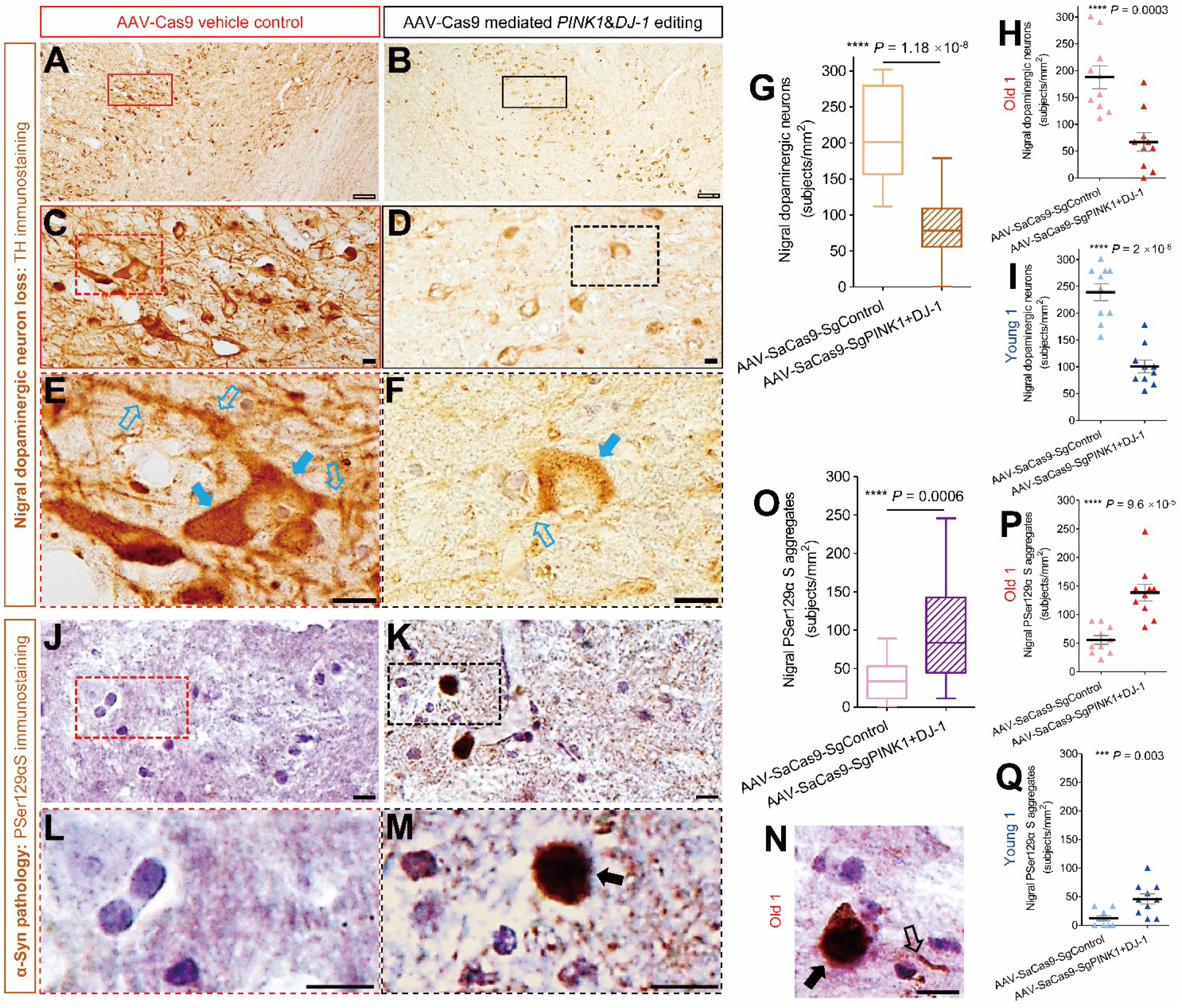
Pathological hallmarks of PD were identified in the SN region of *PINK1* and *DJ-1* edited monkeys. (**A**) TH immunostaining image under low resolution (4 ×) in viral vehicle control side of monkey Old 1’s SN. No morphological changes were found. (**B**) TH immunostaining image under low resolution (4 ×) in *PINK1* & *DJ-1* editing side of monkey Old 1’s SN, which became pale and weak. (**C**) TH immunostaining image under moderate resolution (20 ×) in viral vehicle control side of monkey Old 1’s SN. Some typical dopaminergic neurons were clearly stained, and no obvious pathologic changes were found. (**D**) TH immunostaining image under moderate resolution (20 ×) in *PINK1* & *DJ-1* editing side of monkey Old 1’s SN. Many deformed and lightly stained dopaminergic neurons were identified, showing weak-stained somata and almost completely lost neurites. (**E**) TH immunostaining image under high resolution (40 ×) in viral vehicle control side of monkey Old 1’s SN. A typical dopaminergic neuron was clearly stained with intact soma (solid blue arrows) and branched processes (hollow blue arrows). (**F**) TH immunostaining image under high resolution (40 ×) in *PINK1* & *DJ-1* editing side of monkey Old 1’s SN. A weak-stained dopaminergic neuron was found with incomplete soma boundary (solid blue arrow) and residual process (hollow blue arrow). (**G**) Over 60% of nigral dopaminergic neurons were significantly lost in *PINK1* & *DJ-1* edited side compared with the viral vehicle control side (average of monkey Old 1 and monkey Young 1). (**H-I**) Further quantifications indicated that the nigral dopaminergic neuron loss was also significantly in *PINK1* & *DJ-1* edited side compared with the viral vehicle control side either in monkey Old 1 (**H**) or monkey Young 1 (**I**). (**J**) PSer129αS immunostaining image under high resolution (40 ×) in viral vehicle control side of monkey Old 1’s SN and no typical PSer129αS aggregates were found. (**K**) PSer129αS immunostaining image under high resolution (40 ×) in *PINK1* & *DJ-1* edited side of monkey Old 1’s SN. Some obvious PSer129αS aggregates were identified. (**L**) Enlarged image of the selected area in figure (**J**) indicated that only clearly stained nuclei were found. (**M**) Enlarged image of the selected area in figure (**K**) indicated that a typical PSer129αS aggregate was darkly stained (solid black arrow). (**N**) Other typical examples of PSer129αS aggregates (solid black arrow) and PSer129αS deposit in cell process (hollow black arrow) were found in *PINK1* & *DJ-1* edited side of monkey Old 1’s SN. (**O**) PSer129αS aggregates were significantly more in *PINK1* & *DJ-1* edited side than those in the viral vehicle control side (average of monkey Old 1 and monkey Young 1). (**P-Q**) Further quantifications indicated that the PSer129αS aggregates were also significantly more in *PINK1* & *DJ-1* edited side than those in the viral vehicle control side either in monkey Old 1 (**P**) or monkey Young 1 (**Q**). Scale bar: hollow black: 100 μm; solid black: 10 μm. Data in (G) and (O) are median with minimum to maximum, whereas in (H-I) and (P-Q) are means ± SEM.

Another important PD hallmark, Lewy body pathology, which appears in 77% to 95% PD patients according to different studies (3, 4), is commonly illustrated by the phosphorylated-serine129 α-synuclein (PSer129αS) immunostaining. Although classical Lewy bodies were not identified, the α-synuclein pathology, which is also termed PSer129αS aggregates, was revealed by this staining. The α-synuclein pathology could be considered as an early stage of the Lewy pathology for the following reasons: 1) aggregated α-synuclein is the major component of Lewy bodies and Lewy neurites (34, 35), and about 90% of the α-synuclein aggregates consist of PSer129αS (36); 2) the LBs are probably trigged by abnormal accumulation of PSer129αS in the cytoplasm (37). Illustrated by antibody *ab59264*, one of the most commonly used antibodies for PSer129αS immunostaining (38, 39), some typical PSer129αS aggregates in the cytoplasm were found in SNpc of the genetically edited SN side (**Fig. 4J-K**). Enlarged images showed morphological details of the PSer129αS aggregates (**Fig. 4M-N,** solid black arrow), and the PSer129αS aggregates in cell process was also found (**Fig. 4N,** hollow black arrow), whereas only the nucleus were stained in the AAV-vehicle control side (**Fig. 4L**). Quantitative analysis revealed that the number of PSer129αS aggregates identified in the genetically edited SN side was more than 2 times greater (average of the two monkeys) compared with those in the AAV-vehicle control side (**Fig. 2O**), suggesting that the α-synuclein pathology was successfully identified. Further assessments indicated that the α-synuclein pathology in the genetically edited SN side was significantly more than the AAV-vehicle control side both in monkey Old 1 (**Fig. 4P**) and Young 1 (**Fig. 4Q**) as well.

Collectively, the classic pathology hallmarks of PD were all identified by our current gene co-editing strategy, which could also be attributed to the accurate location of monkey’s SN based on MRI scanning, and it was confirmed by the tracks of viral injections in the brain tissue during the sectioning (**Supplementary Fig. S5A-B**).

### Viral transfection, gene editing consequence and glia roles

To prove that gene editing was effective *in vivo*, we found the expression of Paris/ZNF746, a critical substrate of the PINK1/Parkin signal pathway, was significantly increased in the SN region of an AAV-CRISPR/Cas9 gene edited monkey (**Fig. 5A**), as measured by the Paris level in GFP-positive neurons compared with adjacent GFP-negative neurons. These results not only revealed that the AAVs successfully transfected the nigral neurons, but also caused functional loss of PINK1/Parkin pathway, leading to the up-regulation of Paris, which was consistent with that in the human PD brain (40). Meanwhile, we tested the expression of DJ-1 protein in GFP-positive neurons compared with adjacent GFP-negative neurons in the gene-edited SN, indicating a significant down-regulation of DJ-1 *in vivo* (**Fig. 5B**), which may also contribute to PD pathogenesis. Collectively, the loss-of-function mutations in *PINK1* and *DJ-1* genes were all induced by the nigral AAVs-delivered CRISPR/Cas9 editing, leading to PD pathogenesis.

**Figure 5.**
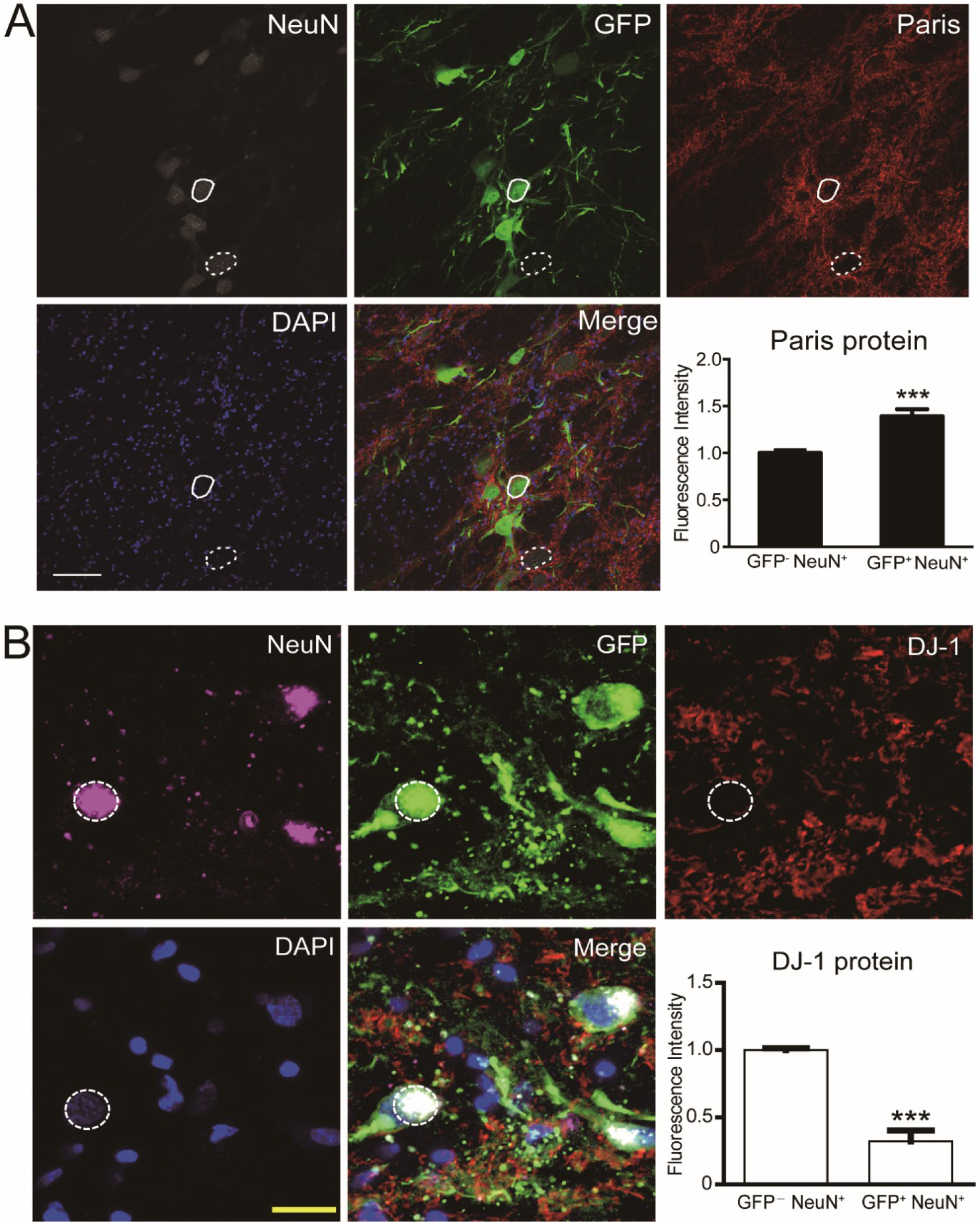
Immunostaining of the functional markers including Paris for *PINK1* and *DJ-1* protein. (**A**) Fluorescence intensity of Paris (red) obtained from GFP-negative or -positive cells was quantified and averaged (GFP negative, n = 18; GFP positive, n = 30). Fluorescence intensity of Paris was increased in GFP-positive cells (*** P < 0.001). (**B**) Fluorescence intensity of DJ-1 (red) obtained from GFP-negative or -positive cells was quantified and averaged (GFP negative, n = 24; GFP positive, n = 24). Fluorescence intensity of DJ-1 was decreased in GFP-positive cells (*** P < 0.001). Scale bar: white: 100 μm, yellow: 20 μm. Data are means ± SEM.

To exclude the inflammatory effects of glial cells activated by the viral injection, which may also trigger PD pathogenesis (41), we tested whether the nigral microglia and astrocyte were transfected and activated by the virus within GFP-positive signals (**Supplementary Fig. S6A**) compared with the GFP-negative area in the SN (**Supplementary Fig. S6B**). We found neither the GFAP-positive cell nor the Iba1-positive cell was co-expressed with GFP (**Supplementary Fig. S6A**), indicating the glial cells were not transfected by the virus. Moreover, no obvious morphological changes of glial cells were found (**Supplementary Fig. S6A-B**). Quantitative assay indicated that Iba1 or GFAP-positive cell count in the GFP-positive area of SN was not significantly increased compared with that in the GFP-negative area (**Supplementary Fig. S6C-D**), indicating the glial cells were not involved in PD pathogenesis during AAV-delivered CRISPR/Cas9 editing procedure.

## Discussion

### Conclusion

In this study, an AAV-delivered CRISPR/Cas 9 system was used to edit the *PINK1* and *DJ-1* genes in the SNs of adult rhesus monkeys to investigate the possible causal relationship between these gene mutations and PD development. After the manipulation, we observed nearly all clinical PD symptoms, including core diagnostic symptoms, such as tremor, bradykinesia, gait abnormality, dramatic changes in hand preference, and rotation induced by the Apo administration, supporting the serious dysfunction of the SN-striatum dopaminergic system. The PD pathological hallmarks were all induced, including loss of more than 60% of dopaminergic neurons in the gene edited SN, pathological α-Syn aggregations and early stage of LBs (Old 1) in the SN. Collectively, these data indicate that core PD phenotypes were all induced by the AAV-delivered CRISPR/Cas9 protocol, and strongly suggest the successful development of a new etiological PD monkey model.

### Technical advantages

Previous studies using viral-based transgenics in adult non-human primate brains induced some PD pathological changes but lacked the core clinical symptoms and LB-like pathology (12, 13). To solve these issues, here we implemented two important technical improvements.

First, ideal PD models should result in more than 50% dopaminergic neuronal loss in the SN (22). To achieve this level of cell death, the AAV could be delivered to cover the whole SN. As the SN is thin, flat, and located deeply within the ventral midbrain, it is difficult to deliver the virus to the SN through normal vertical infusion routes. Here, the unique oblique injection routes (**Fig. 1F**) were employed based on our previous study (27). To cover the whole SN, we injected virus along three parallel paths through the SN, with an average of six injection sites in each path. Therefore, we generated an array of 18 viral injection points across each side of SN (**Fig. 1G**). To ensure precise location of the injection pathways, an accurate MRI-based brain structure localization procedure developed in our lab previously (26) was used and targeted the SN with an error of less than 1 mm, which was confirmed by anatomic examination following monkey sacrifice (**Supplementary Fig. S5**). Actual injection sites and routes are shown in **Supplementary Fig. S4 A-B** and the SN coordinates of each monkey obtained by the MRI scans and calculations are listed in **Supplementary Table S2**.

Second, unlike previous studies that usually manipulate one gene, we edited both the *PINK1* and *DJ-1* genes at the same time via the AAV-delivered CRISPR/Cas9 protocol. This *“dual gene editing”* strategy effectively enhanced PD pathogenesis in the adult rhesus monkeys. This is supported by our preliminary experiments (**Supplementary Table S1**), in which the mutation of either *PINK1* or *DJ-1* alone caused only about 40% of dopaminergic neuronal loss (**Supplementary Fig. S1**), and thus monkeys presented no clear PD symptoms.

### Potential implications

We successfully developed a gene-editing PD etiological monkey model, in which core PD phenotypes were all induced by using an AAV-delivered system to specifically edit the *PINK1* and *DJ-1* genes in the SN region of adult rhesus monkey brain. This work paves an important path for the exploring of specific genetic mechanisms of PD in non-human primates and the screening of biomarkers of PD early diagnosis, as well as for studying of the interactions between environmental risk and genetic mutations and the developing of gene therapies and cell replacement treatments for PD.

## Methods

### Animal ethics

All experiment protocols and animal welfare were approved (*No. IACUC15001*) by the Ethics Committee of Kunming Primate Research Center (AAALAC accredited) and Kunming Institute of Zoology, Chinese Academy of Sciences. All animals involved in this study were treated also in accordance with the National Institute of Health (U.S.A.) Guide for the Care and Use of Laboratory Animals [*8th edition. Washington (DC): National Academies Press; 2011*] and the ARRIVE guidelines for reporting animal research (42). A self-controlled design was used and totally 8 rhesus monkeys (*M. mulatta*) were involved in the study. Four monkeys were involved in the preliminary study for editing *PINK1* or *DJ-1* gene alone, and other four monkeys were involved in this study for co-editing *PINK1* plus *DJ-1* genes together. Detailed information for all the monkeys was listed in **Supplementary Table S1**. Monkeys were cared for by the veterinarians and feeding under standard procedure in individual cage with a standardized light/dark cycle.

Best efforts were made to minimize the number of monkeys involved in this study and their suffering. When the study was finished, the monkeys were anaesthetized and sacrificed for pathological tests. The general condition of monkey Old 1 (94089) became poor at the 14-week mark after viral injections and it was sacrificed for the pathological tests. The monkey Yong 1 (070229) and Young 2 (070209) were selected for pathological examinations at the 20-week mark after viral injection. The remaining monkey (Old 2: 97093) was further observed.

### Viral vector design

The AAV9-SaCas9 vector was obtained from Addgene (#61593). The TBG promoter was replaced with human synapsin promoter for better expression in the brain. The sgRNAs were annealed from DNA oligos with BsaI digest ends. After annealing, the sgRNA segments were inserted into the BsaI site of the vector. The viral titers: vehicle-control: 4.00 × 10^12^; *PINK1*-A: 7.00 × 10^12^; *PINK1*-B: 9.00 × 10^12^; *DJ-1-*A: 1.20 × 10^13^; *DJ-1-*B: 8.00 × 10^12^.

### Mutation efficiency detection in COS7 cell line

The COS7 cell line was cultured by DMEM (high glucose) with 10% fetal bovine serum in 37℃, 5% CO2, and transmitted every 3-4 days. In order to detect the efficiency of sgRNAs, sgRNA and anti-puromycin vectors were transfected in COS7, cells were planted in 6-well plate before transfection, when the coverage rate reached 80%, 6 mg plasmid (4 mg sgRNA vector, 2 mg anti-puro vector) with 4 mg lipofectamine 3000 was added in each well, after 48 hours, puromycin was added to make the final concentration in the medium reached 10 mg/ml to screen out the negative cells. Surviving cells were collected after 24 hours, then digested in lysis buffer with 0.4 mg/ml protease K, the genome was extracted by phenol chloroform. The regions including sgRNA target sites were amplified by PCR, then the PCR products were ligated to T vector by T4 ligase after purification. By transforming competent E.Coli, positive monoclones were selected for sequencing. The results were compared with wild type control sequencing results to determine the efficiency of sgRNA.

Primers used in the PCR amplification included:

*PINK1*-A forward: 5’-CCTGATCTTACCCACTTGC-3’
*PINK1*-A reveres: 5’-GCTCCTGCTCTTCTCCTG-3’
*PINK1*-B forward: 5’-TCACCTTGGCATCTCCTC-3’
*PINK1*-B reveres: 5’-GCTCTACCCGTGCATTTC-3’
*DJ-1*-A forward: 5’-ACAGGTTAATTGCGAAGG-3’
*DJ-1*-A reveres: 5’-GTGGGAAGATGGTTTGAG-3’
*DJ-1*-B forward: 5’-CAGGAAGGAGATTATACTACC-3’
*DJ-1*-B reveres: 5’-CAATAGAACACAAGCAGATG-3’

### Surgeries

#### 1) MRI-guided substantia nigra (SN) localization

Before the anesthesia, all monkeys were injected with atropine (0.5 mg/mL, 1 mL, i.m., Handu, Xinzheng, China). After 10 min, the monkeys were first anesthetized with ketamine (1.0–1.5 mL, 0.1 g/2 mL, i.m., Zhongmu, Taizhou, China) and then injected with pentobarbital sodium (40 mg/mL, 20 mg/kg, i.m., Fluka, Germany) before the surgeries.

During the MRI-guided SN localization procedure, we required a marker that could be clearly seen and fixed without any shift during MRI scanning (Siemens, 3T, Germany) as a reference to locate and calculate the accurate coordinates of the SN. The markers used here were two glass tubes filled with glycerin (3 cm long, internal diameter 0.5 mm, external diameter 0.9 mm), which appeared white with high signal intensity in the MRI T2 images (**Fig. 1E**, white arrows), being fixed onto the skull of each monkey before MRI scanning. Details of the marker fixation and MRI scanning procedures for deep brain region localization have been published previously by our lab (error < 1mm) (26), and this method was successfully used to accurately locate the SN of the rhesus monkey brain in our previous PD study (27). The markers appeared as two light and parallel lines in the middle coronal section during MRI scanning; as a result, this coronal plane was accurately determined by the two markers with a definitive locations: 10 mm before the **A** (anterior) / **P** (posterior) **Zero** (midpoint of left and right ears of the monkey), 5 mm to left and right of the midsagittal plane respectively, and about 1 mm deep in the skull. This coordinate was chosen from the Rhesus Monkey Brain Atlas: *Paxinos G, Toga AW, Huang X-F (1999), The Rhesus Monkey Brain in Stereotaxic Coordinates: Academic Pr. 163 p.*, which was used as a reference to locate the SN.

The SN was clearly seen on the T2 images with low signal intensity, appearing black (**Fig. 1F**, green circle). The optimum section of SN acquired from T2 MRI scanning for the calculations of coordination guiding for viral injections was selected based on the SN structure characteristics and imaging quality of MRI. Anatomically, the clear morphology and largest area is nearly located in the middle from the anterior to the posterior of the SN. The MRI image quality depends on the fine definition and high signal to noise ratio. According to the above standards, representative images of MRI-located SN in monkey Old 1 along the coronal plane and calculations for the path of viral injections are shown in **Fig. 1F**. The symmetry axis of the monkey’s brain in this coronal plane was labeled with a blue line and the path targeted along the SN was labeled with a white line (the length from endpoint of the SN to the skull is represented by a “d”). This was intersected with the skull and the crossover point was labeled with a red point, with a vertical line from this red point to the blue line then indicated with a red line, and the corresponding length represented by an “s”. Therefore, a right-angled triangle was formed and the angle θ was calculated by ****sin θ = s / d**** with the lengths of “s” and “d” being measured by MRI software. As a result, the key parameters used for viral injection were calculated and are listed in **Supplementary Table S2**. As described above, “s” is the distance to the midsagittal plane (mm), “d” is the distance from the skull to the SN endpoint (mm), and “θ” is the angle from the gray line to the blue line (°). Based on the coordinates, the injection path (white line) was determined for the viral injections. For each monkey, when the MRI scans were completed, the markers were removed from the skull and the wound was closed. After that, each monkey was given penicillin (800 KU, i.m., H13021634, Shiyao, Shijiazhuang, China) daily for one week.

#### 2) Viral injections

After one week of recovery, each monkey underwent another operation for the viral injections and the anesthesia was same as above. The injection route of each monkey was decided by the coordinates listed in **Supplementary Table S2**, and the corresponding location on each monkey’s skull (red point) was drilled accordingly. When the virus-filled micro-injector arrived at the endpoint of the SN, a qualified injection strategy was adopted to cover the whole SN with multi-points. On average, six points along each route were arranged for viral injections (50 μL in total) in every monkey, with a separation distance of 1 mm along the path of the SN and the total length of the path was about 6 mm. Of note, the SN region is not limited to one coronal plane in MRI scanning. We added two other routes with the same parameters in two coronal plans before and after the optimum SN plane with an interval of 2 mm between each, thus a total of three parallel injection paths (150 μL in total) and 18 injection points were involved in the viral injections (**Fig. 1G**). The viral titers were 10^12^ or 10^13^. After each point was injected, the micro-injector was lifted by 1 mm for the next injection. The administration speed was held constant at 800 nL/min by an injection pump (World Precision Instruments, Inc., UMP 3-4, USA), and the time interval between each injection path was 5 min (43). After the injections, the wound was sutured. All monkeys were administrated with penicillin (800 KU, i.m., H13021634, Shiyao, Shijiazhuang, China) daily for one week.

### Behavioral data collection

To evaluate the Parkinsonian behavior of monkeys and the SN-striatum pathway dysfunctions, we collected and analyzed the behavioral data from the following three aspects: **1)** quantification of the PD symptoms by videotaping the monkey’s daily activities and then scoring it using the improved *Kurlan* scale (a monkey-parkinsonism rating scale); **2)** the SN-striatum pathway dysfunctions by using apomorphine (Apo)-induced rotation test; and **3)** the refined finger movement deficits by the food grasping task.

**1)** To identify and quantify the PD symptoms on gene edited monkeys, video recordings were essential for behavioral data collection. The video recordings of the monkey’s behavior were collected by digital cameras (Sony HDR-XR260, Japan). Briefly, each single-caged monkey was recorded without external disturbance for 1 h by the camera located 1 m in front of the cage at 10:00-11:00 am. After the behavioral data collection, the 1 h video was used for the behavioral assessment.

The next step of video analyzing process was based on previous studies (27, 44). Here, two well trained observers were involved in the video clips analyzing. The videos were randomly labeled by the experimenter with the letters from A to D. Blind to conditions or treatments of the monkey, two observers individually assigned the video clips to score separately using the improved *Kurlan* scale.

The improved *Kurlan* scale has been widely accepted as a standard scale for PD research on Old World Monkeys (29). Briefly, this scale includes 4 parts: part A: Parkinsonian features; part B: drug-related side effects; part C: overall level of activity; part D: clinical staging (28). The PD score is calculated by using the part A of this scale, which includes seven PD measurements: tremor, posture, gait, bradykinesia, balance, gross motor skill, and defense reaction (28). According to the part A of this scale with a total PD score of 20, the score of each item was evaluated and assigned by the severity (*tremor: 0~3; posture: 0~2; gait: 0~4; bradykinesia: 0~4; balance: 0~2; gross motor skill: 0~3; defense reaction: 0~2*). The total PD score was calculated with the sum of each score evaluated from the individual items. Moreover, no drug-related side effects were observed according to the part B (dyskinesias, vomiting, and psychological disturbance) of this scale, and the overall level of activity of the monkeys was “moderate hypokinesia: sparse movements” according to the part C of this scale. The “Clinical staging” of the monkeys was evaluated by the part D of the scale mentioned above (stage II). The standards refer to two behavioral assessments: **a)** hemi/bilateral Parkinsonism evaluated by the behavioral performances and pathological staining; **b)** does not use affected upper limb for walking.

After the scoring, if there were no obvious differences (less than two scores) between these two observers in evaluating each item of the scale, the data was pooled together for creating figures and performing statistics.

**2)** To examine the dysfunction of the SN-striatum dopaminergic system by CRISPR/Cas9-*PINK1* and *DJ-1* editing, we used a classic pharmacological test with Apo administration. This rotation test consisted of two steps: first, the spontaneous rotations of monkeys to the lesioned side were counted in a 30-min section of a video episode, which had collected by the protocol mentioned above. The duration of each collected video was 1 h, but a 30-min section was selected for counting the rotations in comparison to the later Apo-induced rotations. Second, for the Apo-induced rotations to the healthy side, a new video was recorded immediately after the monkeys were injected with Apo [R-(-)-apomorphine hydro-chloride hemihydrate, A4393, Sigma,]. Evaluation was also based on a 30-min section, which began 15 min after the Apo injection and ended at the 45 min accordingly. For each test, Apo (2 mg) was dissolved (1 mg/mL; 2 mL, i.m.) in vitamin C solution (0.5 mg/2 mL) 5 min before the injection. After that, all the videos were randomly labeled by the experimenter with the numbers from 1 to 8. Blind to conditions or treatments of the monkey, two observers individually counted the rotations from all collected above videos.

**3)** To test refined movement deficits of each monkey during PD progression, a food grasping task was employed (45). During the task, each individually caged monkey was trained to fetch a piece of food as a reward from a food box in front of its cage. After the monkey focused on the task and fetching behavior stabilized, baseline data were collected. The time (latency) and accuracy (%) of food grasping were calculated every 10 trails, and the hand preference was also recorded. Each monkey had to finish 180 trails over three successive days (60 trails per day) of each testing week. Peanuts, diced apples, or food pellets were selected as the rewards depending on each monkey’s preference.

### Pathology

#### 1) bran tissue sectioning and anatomic location of substantia nigra (SN)

After the behavioral data collection, three monkeys (Young 1, Young 2, and Old 1) were first euthanized with ketamine (2 mL, 0.1 g/2 mL, i.m. Zhongmu, Taizhou, China) and then administrated with pentobarbital sodium (40 mg/mL, 20 mg/kg, i.m. Fluka, Germany). After the anesthesia, perfusions were carried out. The monkeys’ brains were acquired and fixed in 4% paraformaldehyde-0.01 M PBS for one week (Young 1 and Old 1) and then gradually equilibrated with 20% and 30% sucrose. Coronal sections of the whole SN of Young 1 and Old 1 were obtained by sectioning the tissue (20 μm) on a freezing microtome (Leica, CM1850UV-1-1, Germany) at −24 °C.

The main brain structure involved in PD pathology is the SN, a paired flat shaped structure located deep in the narrow mesencephalon (27). The SN was located from −9.90 mm to −17.78 mm AP from bregma according to the monkey brain atlases *“The Rhesus Monkey Brain in Stereotaxic Coordinates, George Praxinos et al., Academic Press, 1999”*, and it was further confirmed by the MRI data, proving the length of the SN is about 8.00 mm from anterior to the posterior.

Based on the studies by Chu et al. (46–48) and the neuropathological assessment for PD by Dickson et al. (49), the pathological criteria for selecting the optimum section of SN for pathologic validation was adopted. It was in the ventral midbrain and defined ventrally by the cerebral peduncle and medially by the third cranial nerve (3n) rootlets in coronal section. Here, the length of the 3n combined with clear SN in the monkey brain atlas was 2.25 mm (−10.80 mm to −13.05 mm AP from bregma), about 30% length of the whole SN (2.25/8.00 = 28.13%).

Under the direction of the anatomical guidance, the bilateral side of whole SN from the monkeys Yong 1 and Old 1 were cut into 80 coronal sections by a freezing microtome. The section thickness was 20 μm, the interval between sections was 80 μm, and therefore, the 80 sections basically covered the entire SN: 80 sections × (20 + 80) μm = 8 mm. The optimum sections for the dopaminergic neuron staining and counting were ~20 coronal sections (80 sections × 28.55% = 22 sections).

#### 2) staining

In monkeys Young 1 and Old 1, sections including the target brain areas were selected and mounted onto slides for staining based on our previous protocols (44, 47). In TH staining, sections of the SN were treated with 3% H_2_O_2_ (5 min, Maixin, Fuzhou, China), 3% triton X-100 (4 min, Solarbio, Beijing, China), and goat serum (15 min, Maixin, Fuzhou, China) and then incubated with rabbit anti-TH antibody (1: 1,000, AB152, Millipore, USA) overnight at 4 °C. The sections were then incubated with secondary antibodies (PV-9000, 30 min, ZSGB-BIO, Beijing, China) at 37 °C. Afterwards, sections were interacted with DAB (DAB-1031Kit, 20 ×, Maixin, Fuzhou, China) and counterstained with hematoxylin.

Previously published protocols were followed in the staining of PSer129αS (50, 51). The sections were firstly treated with proteinase K (Aladdin, Shanghai, China) solution at 37 °C for 30 min: 50 μg/mL proteinase K was dissolved in the buffer containing 10 mM Tris-HCl, pH = 7.8, 100 mM NaCl, and 0.1 % Tween-20 (Bio-Rad, USA). The sections were then subsequently treated as same as the TH staining, except the antibody incubation with rabbit anti- PSer129αS antibody (1: 100, ab59264, Abcam, UK) overnight at 4 °C.

After the immunostaining, all sections were dehydrated with the alcohol gradients and made transparent with xylene. Neutral balsam was used to cover the slides and digital pictures were collected on a microscope (Olympus, CX41; camera: Olympus DP25; software: CellSens Entry 1.4.1; Japan). Ten sections containing both the left and right SN region from each monkey were selected and used for counting of TH positive cells and PSer129αS aggregates.

In Young 2, the brain was dissected, cut into small blocks after perfusion, fixed with 4% PFA in phosphate buffer, and then equilibrated in 30% sucrose. Fixed and equilibrated brain tissue blocks were cut into 50 μm cortical sections with a Microm HM525 cryostat. Sections were washed for 5 min in 0.01 M PBS containing 5% bovine serum albumin (BSA) and 0.3% Triton X-100, and then incubated with primary antibodies (in 0.01 M PBS with 1% BSA and 0.3% Triton X-100) overnight at 4 °C and subsequently with corresponding secondary antibodies (1:1,000, Alexa-Fluor-conjugated, Invitrogen). DAPI was used to label the nuclei and sections were mounted with 75% glycerol. Primary antibodies used were included: GFP antibody (ab6673, Abcam), NeuN antibody (ABN78, Millipore), Paris antibody (75-195, NeuroMab), DJ-1 antibody (ab201147, Abcam), another NeuN antibody (MABN377, Millipore), GFAP antibody (SMI-21R, Covance Research Products Inc), Iba 1 antibody (019-19741, Wako), and another GFP antibody (ab13970, Abcam).

#### 3) stereological counting

The substantia nigra pars compacta (SNpc) was selected for cell counting because most nigral dopamine cells are in this area. The cells were counted by ImageJ software (National Institutes of Health, Bethesda, Maryland, USA) under a 40 × objective (counting frame) to correctly recognize the morphology of individual cell. Each selected counting area of every section from vehicle control side and the viral-delivered gene editing side was similar and unbiased located around the middle of the SNpc long axis and further determined by the morphological characters, including the anatomical appearance: ellipsoid shaped and defined ventrally by the cerebral peduncle and medially by the third cranial nerve (3n), and the cellular properties: spindle-shaped, pigmented and well evenly distributed. These standardized methods proposed for assessment of neuronal loss in the SNpc was adopted from the neuropathological assessment for PD by Dickson et al. (49) and related studies (52, 53).

Nigral TH^+^ cell count by using ImageJ software on each side of the SN for the two monkeys were performed totally in 20 sections (10 sections per monkey). Data obtained from the vehicle control side of the SN in each monkey including 20 sections in total were pooled together and averaged. Accordingly, for the viral-delivered gene editing side of SN, cell counting from 20 sections was also averaged. Each TH+ cell selected by the imageJ software must be confirmed again with intact cellular structure and nucleus. For counting the aggregates of PSer129αS, the method was similar.

#### 4) Confocal imaging and cell quantification

Confocal z-stack images were acquired on a Nikon A1 confocal laser microscope system. NIH ImageJ software was used to analyze the mean fluorescence intensity of immunoreactive cells. The fluorescence intensity of Paris from GFP^+^ NeuN^+^ and GFP^−^ NeuN^−^ nigral cells were analyzed to quantify the expression level of PINK1, and the fluorescence intensity of DJ-1 from GFP+ NeuN+ and GFP-NeuN-nigral cells were also analyzed to quantify the expression level of DJ-1.

### Statistical analysis

Statistical analysis was performed with SPSS software (IBM SPSS Statistics 25, free version). Correlations were performed with SPSS 25 (regression-curve fit). ANOVA (homogeneous variance) or NPar tests: Mann-Whitney Test (non-homogeneous variance) were used to compare the means/distributions of PD scores, latency, accuracy, rotations, TH^+^ cell count, Iba 1^+^ cell count, GFAP+ cell count, PSer129αS aggregates, and fluorescence intensity of DJ-1 in GFP-positive or GFP-negative neurons. Two-tailed Student’s *t*-test was used to compare the means/distributions of fluorescence intensity of Paris in GFP positive or GFP-negative neurons. Significance was set as: *\* P < 0.05, ** P < 0.01, *** P < 0.001*.

## Acknowledgements

We would like to thank Mr. Hongwei Li for his hard work on monkey training and video collection, and Miss Ling Zhao for technical support with monkey feeding and housing. This work was supported by the National Key R&D Program of China (2018YFA0801403), the Key Realm R&D Program of Guangdong Province (2019B030335001), the Strategic Priority Research Program of the Chinese Academy of Sciences (XDB32060200), the National Program for Key Basic Research Projects (973 Program: 2015CB755605), National Natural Science Foundation of China (81471312, 81771387, 81460352, 81500983, 31700897, 31700910, 31800901, 31625013, 91732302), the Applied Basic Research Programs of Science and Technology Commission Foundation of Yunnan Province (2017FB109, 2018FB052, 2018FB053, 2019FA007), CAS “Light of West China” Program, Shanghai Brain-Intelligence Project from STCSM (16JC1420501), Shanghai Municipal Science and Technology Major Project (2018SHZDZX05), Open Large Infrastructure Research of Chinese Academy of Sciences, and China Postdoctoral Science Foundation (2018M631105).

## Author contributions

H.L. contributed to experimental design, surgery, behavioral tests, pathology, data analysis, and drafting; S.H.W. contributed to surgery, monkey sacrifice, and figure formatting; X.L. contributed to viral design and off-target tests; X.M. contributed to the immunostainings and data analysis. T.L.C. contributed to cell culture and PCR; Z.F.C. contributed to immunostainings; J.W. contributed to SN location surgeries; L.B.L. contributed to the technical support for NHP research; L.L. contributed to MRI scanning; L.Q.X. contributed to diagnosis of MRI data, W.C.W. contributed to drafting; Y.Z.H. contributed to editing the manuscript, H.S.J. contributed to Paris staining suggestions; Y.Y. contributed to providing funds and comments; Z.L.Q. contributed to providing funds, manuscript drafting, and revision; and X.T.H. contributed to providing funds, the study design, manuscript drafting and editing.

## References

1. Poewe, W, Seppi, K, Tanner, CM, et al. Parkinson disease. Nat Rev Dis Primers. 2017; 3: 17013.

2. Goedert, M, Spillantini, MG, Del Tredici, K, et al. 100 years of Lewy pathology. Nat Rev Neurol. 2013; 9(1): 13–24.

3. Gibb, WR. Idiopathic Parkinson's disease and the Lewy body disorders. Neuropathol Appl Neurobiol. 1986; 12(3): 223–34.

4. Dickson, DW. Neuropathology of Parkinson disease. Parkinsonism Relat Disord. 2018; 46 Suppl 1: S30–S3.

5. Olanow, CW, Kieburtz, K, Schapira, AH. Why have we failed to achieve neuroprotection in Parkinson's disease? Ann Neurol. 2008; 64 Suppl 2: S101–10.

6. Savica, R, Grossardt, BR, Bower, JH, et al. Incidence and pathology of synucleinopathies and tauopathies related to parkinsonism. JAMA Neurol. 2013; 70(7): 859–66.

7. Buffalo, EA, Movshon, JA, Wurtz, RH. From basic brain research to treating human brain disorders. Proc Natl Acad Sci U S A. 2019; 116: 26167–72.

8. Mattison, JA, Vaughan, KL. An overview of nonhuman primates in aging research. Exp Gerontol. 2017; 94: 41–5.

9. Hisahara, S, Shimohama, S. Toxin-induced and genetic animal models of Parkinson's disease. Parkinsons Dis. 2010; 2011: 951709.

10. Dawson, TM, Ko, HS, Dawson, VL. Genetic animal models of Parkinson's disease. Neuron. 2010; 66(5): 646–61.

11. Kalia, LV, Lang, AE. Parkinson's disease. Lancet. 2015; 386(9996): 896–912.

12. Kirik, D, Annett, LE, Burger, C, et al. Nigrostriatal alpha-synucleinopathy induced by viral vector-mediated overexpression of human alpha-synuclein: a new primate model of Parkinson's disease. Proc Natl Acad Sci U S A. 2003; 100(5): 2884–9.

13. Eslamboli, A, Romero-Ramos, M, Burger, C, et al. Long-term consequences of human alpha-synuclein overexpression in the primate ventral midbrain. Brain. 2007; 130(Pt 3): 799–815.

14. Liu, Z, Li, X, Zhang, JT, et al. Autism-like behaviours and germline transmission in transgenic monkeys overexpressing MeCP2. Nature. 2016; 530(7588): 98–102.

15. Chen, Y, Yu, J, Niu, Y, et al. Modeling Rett Syndrome Using TALEN-Edited MECP2 Mutant Cynomolgus Monkeys. Cell. 2017; 169(5): 945–55 e10.

16. Yang, W, Liu, Y, Tu, Z, et al. CRISPR/Cas9-mediated PINK1 deletion leads to neurodegeneration in rhesus monkeys. Cell Res. 2019.

17. Wiedenheft, B, Sternberg, SH, Doudna, JA. RNA-guided genetic silencing systems in bacteria and archaea. Nature. 2012; 482(7385): 331–8.

18. Jinek, M, Chylinski, K, Fonfara, I, et al. A programmable dual-RNA-guided DNA endonuclease in adaptive bacterial immunity. Science. 2012; 337(6096): 816–21.

19. Mali, P, Yang, L, Esvelt, KM, et al. RNA-guided human genome engineering via Cas9. Science. 2013; 339(6121): 823–6.

20. Cong, L, Ran, FA, Cox, D, et al. Multiplex genome engineering using CRISPR/Cas systems. Science. 2013; 339(6121): 819–23.

21. Gasser, T. Molecular pathogenesis of Parkinson disease: insights from genetic studies. Expert Rev Mol Med. 2009; 11: e22.

22. Beal, MF. Experimental models of Parkinson's disease. Nature reviews Neuroscience. 2001; 2(5): 325–34.

23. Moore, DJ, West, AB, Dawson, VL, et al. Molecular pathophysiology of Parkinson's disease. Annu Rev Neurosci. 2005; 28: 57–87.

24. Dolgacheva, LP, Berezhnov, AV, Fedotova, EI, et al. Role of DJ-1 in the mechanism of pathogenesis of Parkinson's disease. J Bioenerg Biomembr. 2019; 51(3): 175–88.

25. Collier, TJ, Kanaan, NM, Kordower, JH. Ageing as a primary risk factor for Parkinson's disease: evidence from studies of non-human primates. Nature reviews Neuroscience. 2011; 12(6): 359–66.

26. Jing, W, Wenchao, W, Lin, L, et al. A new MRI approach for accurately implanting microelectrodes into deep brain structures of the rhesus monkey (Macaca mulatta). J Neurosci Methods. 2010; 193(2): 203–9.

27. Lei, X, Li, H, Huang, B, et al. 1-methyl-4-phenylpyridinium stereotactic infusion completely and specifically ablated the nigrostriatal dopaminergic pathway in rhesus macaque. PLoS One. 2015; 10(5): e0127953.

28. Smith, RD, Zhang, Z, Kurlan, R, et al. Developing a stable bilateral model of parkinsonism in rhesus monkeys. Neuroscience. 1993; 52(1): 7–16.

29. Imbert, C, Bezard, E, Guitraud, S, et al. Comparison of eight clinical rating scales used for the assessment of MPTP-induced parkinsonism in the Macaque monkey. J Neurosci Methods. 2000; 96(1): 71–6.

30. Bezard, E, Przedborski, S. A tale on animal models of Parkinson's disease. Mov Disord. 2011; 26(6): 993–1002.

31. Przedborski, S, Jackson-Lewis, V, Popilskis, S, et al. Unilateral MPTP-induced parkinsonism in monkeys. A quantitative autoradiographic study of dopamine D1 and D2 receptors and re-uptake sites. Neurochirurgie. 1991; 37(6): 377–82.

32. Chitkara, B, Durcan, MJ, Campbell, IC. Apomorphine-induced stereotypy: function of age and rearing environment. Pharmacol Biochem Behav. 1984; 21(4): 671–3.

33. Hirschhorn, ID, Makman, MH, Sharpless, NS. Dopamine receptor sensitivity following nigrostriatal lesion in the aged rat. Brain Res. 1982; 234(2): 357–68.

34. Spillantini, MG, Crowther, RA, Jakes, R, et al. alpha-Synuclein in filamentous inclusions of Lewy bodies from Parkinson's disease and dementia with lewy bodies. Proc Natl Acad Sci U S A. 1998; 95(11): 6469–73.

35. Spillantini, MG, Schmidt, ML, Lee, VM, et al. Alpha-synuclein in Lewy bodies. Nature. 1997; 388(6645): 839–40.

36. Fujiwara, H, Hasegawa, M, Dohmae, N, et al. alpha-Synuclein is phosphorylated in synucleinopathy lesions. Nat Cell Biol. 2002; 4(2): 160–4.

37. Wakabayashi, K, Tanji, K, Mori, F, et al. The Lewy body in Parkinson's disease: molecules implicated in the formation and degradation of alpha-synuclein aggregates. Neuropathology. 2007; 27(5): 494–506.

38. Bae, EJ, Yang, NY, Song, M, et al. Glucocerebrosidase depletion enhances cell-to-cell transmission of alpha-synuclein. Nat Commun. 2014; 5: 4755.

39. Shahmoradian, SH, Lewis, AJ, Genoud, C, et al. Lewy pathology in Parkinson's disease consists of crowded organelles and lipid membranes. Nat Neurosci. 2019; 22(7): 1099–109.

40. Shin, JH, Ko, HS, Kang, H, et al. PARIS (ZNF746) repression of PGC-1alpha contributes to neurodegeneration in Parkinson's disease. Cell. 2011; 144(5): 689–702.

41. Cao, JJ, Li, KS, Shen, YQ. Activated immune cells in Parkinson's disease. J Neuroimmune Pharmacol. 2011; 6(3): 323–9.

42. Kilkenny, C, Browne, WJ, Cuthill, IC, et al. Improving bioscience research reporting: the ARRIVE guidelines for reporting animal research. PLoS Biol. 2010; 8(6): e1000412.

43. Wu, SH, Liao, ZX, J, DR, et al. Comparative study of the transfection efficiency of commonly used viral vectors in rhesus monkey (Macaca mulatta) brains. Zool Res. 2017; 38(2): 88–95.

44. Li, H, Lei, X, Huang, B, et al. A quantitative approach to developing Parkinsonian monkeys (Macaca fascicularis) with intracerebroventricular 1-methyl-4-phenylpyridinium injections. J Neurosci Methods. 2015; 251: 99–107.

45. Courtine, G, Roy, RR, Raven, J, et al. Performance of locomotion and foot grasping following a unilateral thoracic corticospinal tract lesion in monkeys (Macaca mulatta). Brain. 2005; 128(Pt 10): 2338–58.

46. Chu, Y, Kompoliti, K, Cochran, EJ, et al. Age-related decreases in Nurr1 immunoreactivity in the human substantia nigra. J Comp Neurol. 2002; 450(3): 203–14.

47. Chu, Y, Kordower, JH. Age-associated increases of alpha-synuclein in monkeys and humans are associated with nigrostriatal dopamine depletion: Is this the target for Parkinson's disease? Neurobiol Dis. 2007; 25(1): 134–49.

48. Chu, Y, Muller, S, Tavares, A, et al. Intrastriatal alpha-synuclein fibrils in monkeys: spreading, imaging and neuropathological changes. Brain. 2019.

49. Dickson, DW, Braak, H, Duda, JE, et al. Neuropathological assessment of Parkinson's disease: refining the diagnostic criteria. Lancet Neurol. 2009; 8(12): 1150–7.

50. Tanji, K, Mori, F, Mimura, J, et al. Proteinase K-resistant alpha-synuclein is deposited in presynapses in human Lewy body disease and A53T alpha-synuclein transgenic mice. Acta Neuropathol. 2010; 120(2): 145–54.

51. Huang, B, Wu, S, Wang, Z, et al. Phosphorylated alpha-Synuclein Accumulations and Lewy Body-like Pathology Distributed in Parkinson's Disease-Related Brain Areas of Aged Rhesus Monkeys Treated with MPTP. Neuroscience. 2018; 379: 302–15.

52. Kimura, K, Inoue, KI, Kuroiwa, Y, et al. Propagated but Topologically Distributed Forebrain Neurons Expressing Alpha-Synuclein in Aged Macaques. PLoS One. 2016; 11(11): e0166861.

53. Wegrzynowicz, M, Bar-On, D, Calo, L, et al. Depopulation of dense alpha-synuclein aggregates is associated with rescue of dopamine neuron dysfunction and death in a new Parkinson's disease model. Acta Neuropathol. 2019; 138(4): 575–95.

