## Supplementary Figure S1-S6 and Supplementary Table S1-S4 for "Co-editing *PINK1* and *DJ-1* genes via AAV-delivered CRISPR/Cas9 system in adult monkey brains elicits classic Parkinsonian phenotypes"

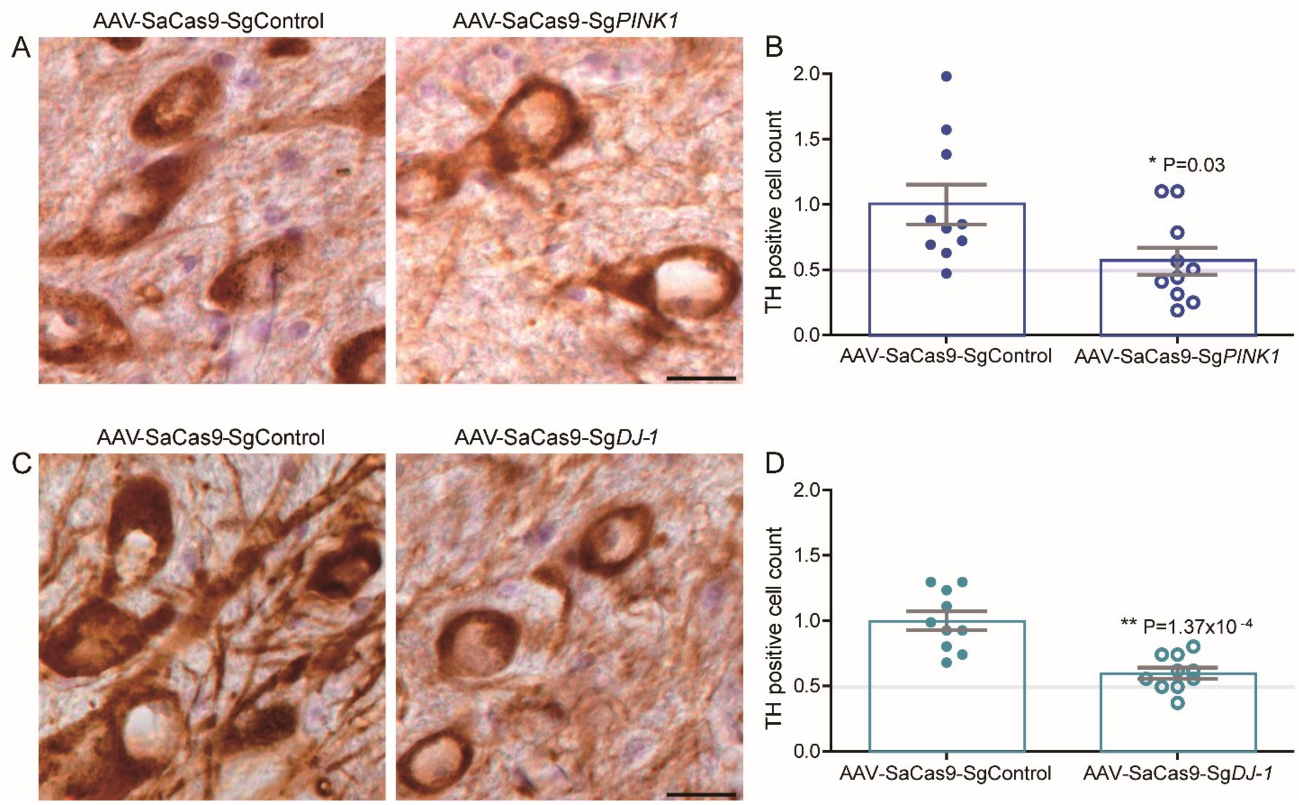


**Fig. S1. Nigral dopaminergic cell loss of *PINK1* or *DJ-1* gene edited alone by AAV-delivered CRISPR/Cas9.** (**A**) TH positive neurons stained in the SN area of *PINK1* gene edited alone by AAV-delivered CRISPR/Cas9 and control side. (**B**) TH positive cells were significantly reduced in the *PINK1* gene edited side but not more than 50% loss. (**C**) TH positive neurons stained in the SN area of *DJ-1* gene edited alone by AAV-delivered CRISPR/Cas9 and control side. (D) TH positive cells were also significantly reduced in the *DJ-1* gene edited side but not more than 50% loss. Scale bar: 20 µm. Data are means ± SEM.

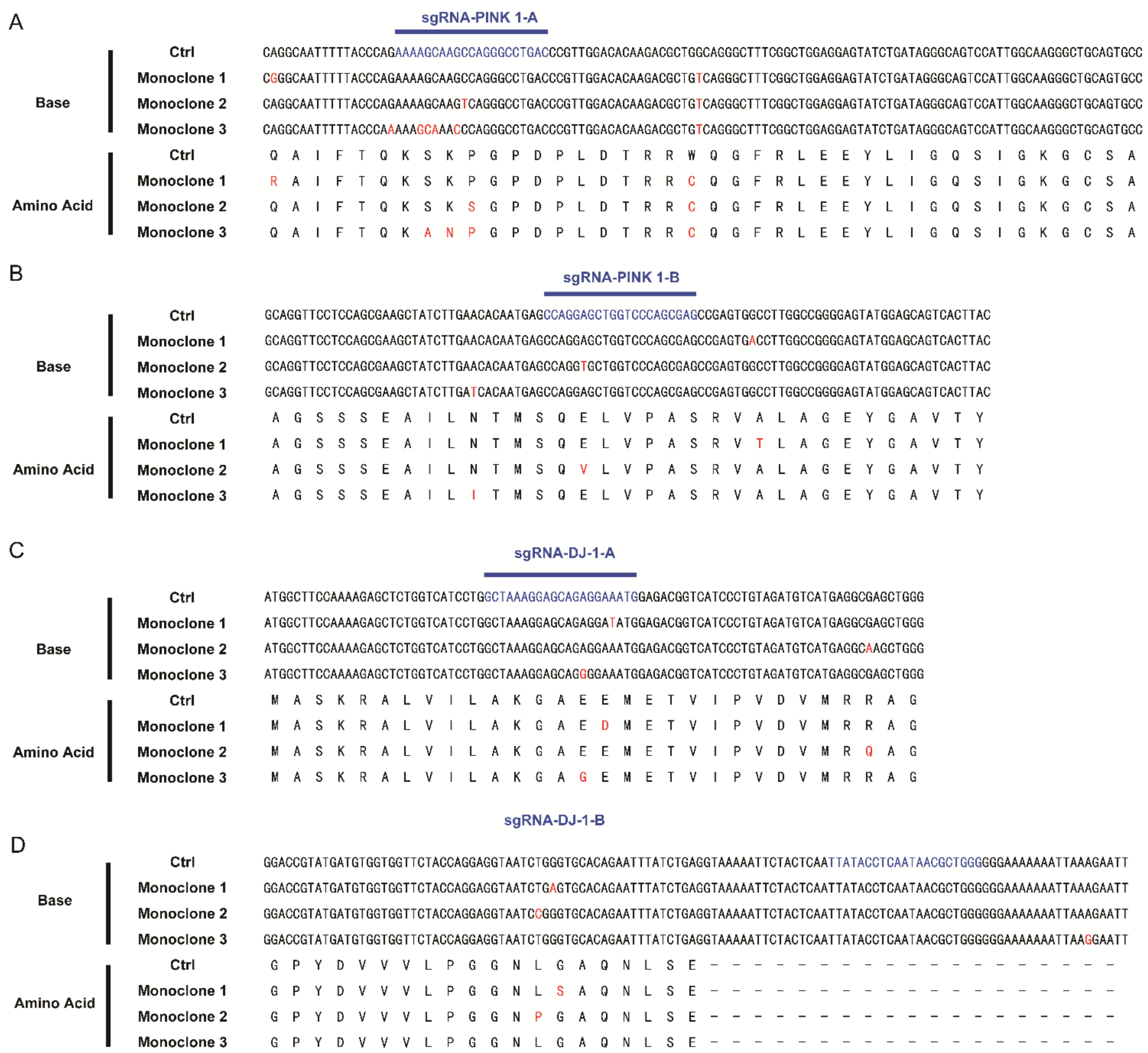

**Fig. S2. Identification of genetic mutations caused by SaCas9 and sgRNAs.** (**A-D**) Examples of monoclone mutations near target site caused by sgRNAs in COS7 cell line. Red highlights show mutated base and amino acid, and blue highlights show sgRNAs target sites.

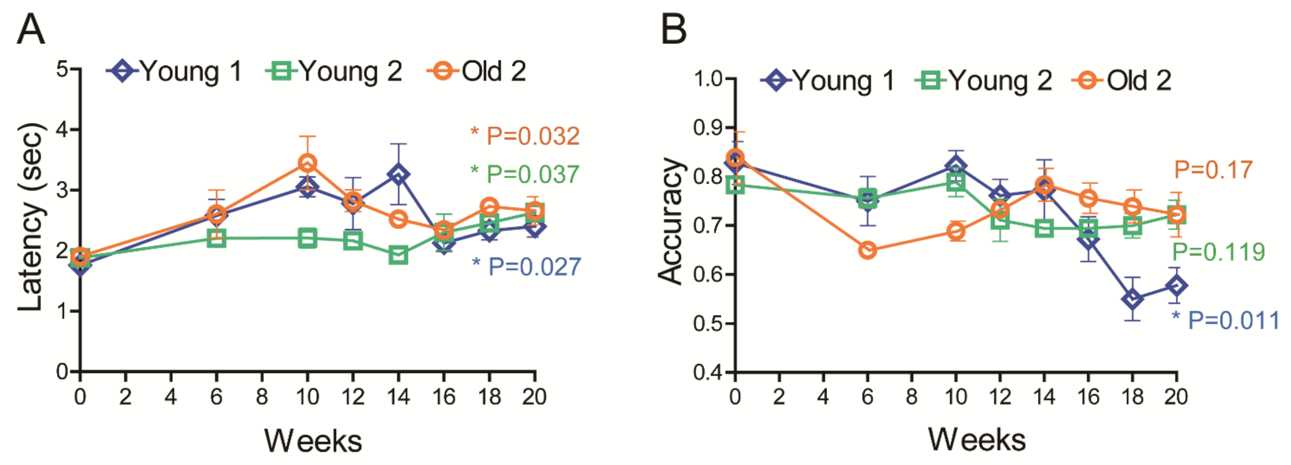


**Fig. S3. Change in latency and accuracy in the three monkeys who altered their hand preference after AAV-delivered CRISPR/Cas9 editing of *PINK1* & *DJ-1* in their SNs.** (**A**) Latency of the three monkeys was significantly increased at 20 weeks after viral injections in comparison to the baseline (0 week). (**B**) Accuracy was significantly decreased in monkey Young 1 at 20 weeks after viral injections in comparison to the baseline (0 week). Data are means ± SEM.

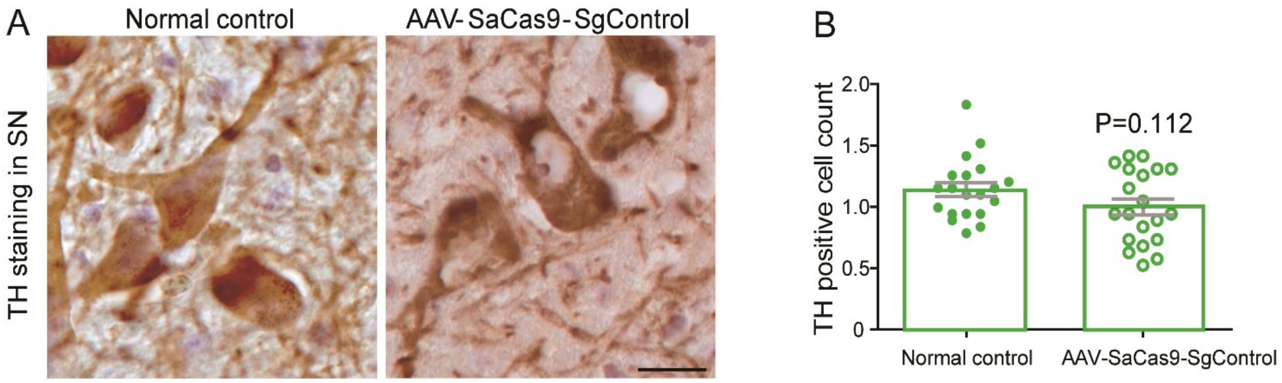


**Fig. S4. Comparison of TH staining in SN region of age-matched normal control monkey and AAV-SaCas9-SgControl.** (**A**) There were no obvious morphological differences in nigral TH-positive cells between these two groups. (**B**) The number of TH-positive cells was not significantly decreased in AAV-SaCas9-SgControl. Data is means ± SEM. Scale bar: 20 μm.

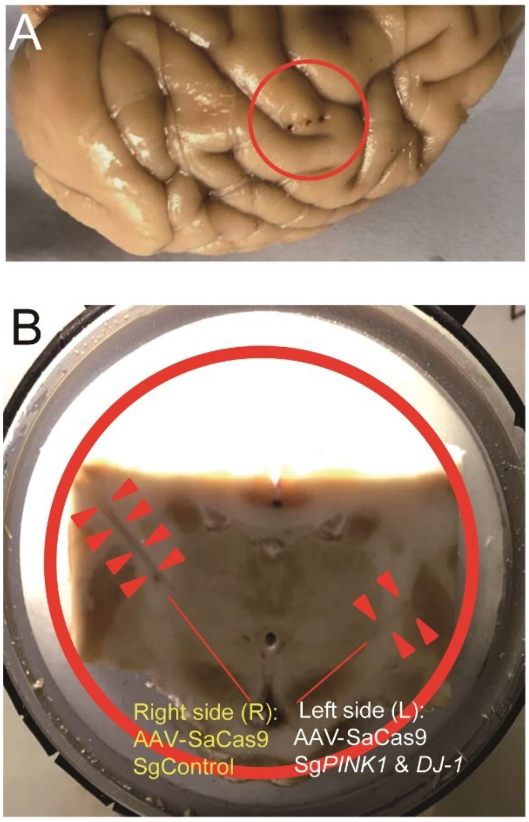


**Fig. S5. Anatomical evidence for the accurate injections of AAV-delivered CRISPR/Cas 9 into the SN.** (**A**) Injection pores enlarged by needles with a diameter of 0.5 mm in the brain of monkey Old 1 (small red circle). (**B**) Red arrows indicate enlarged injection paths directly targeting to the SN regions (red lines). The big red circle indicates the coronal section of brain tissue cut along the middle-injected pore shown in figure A.

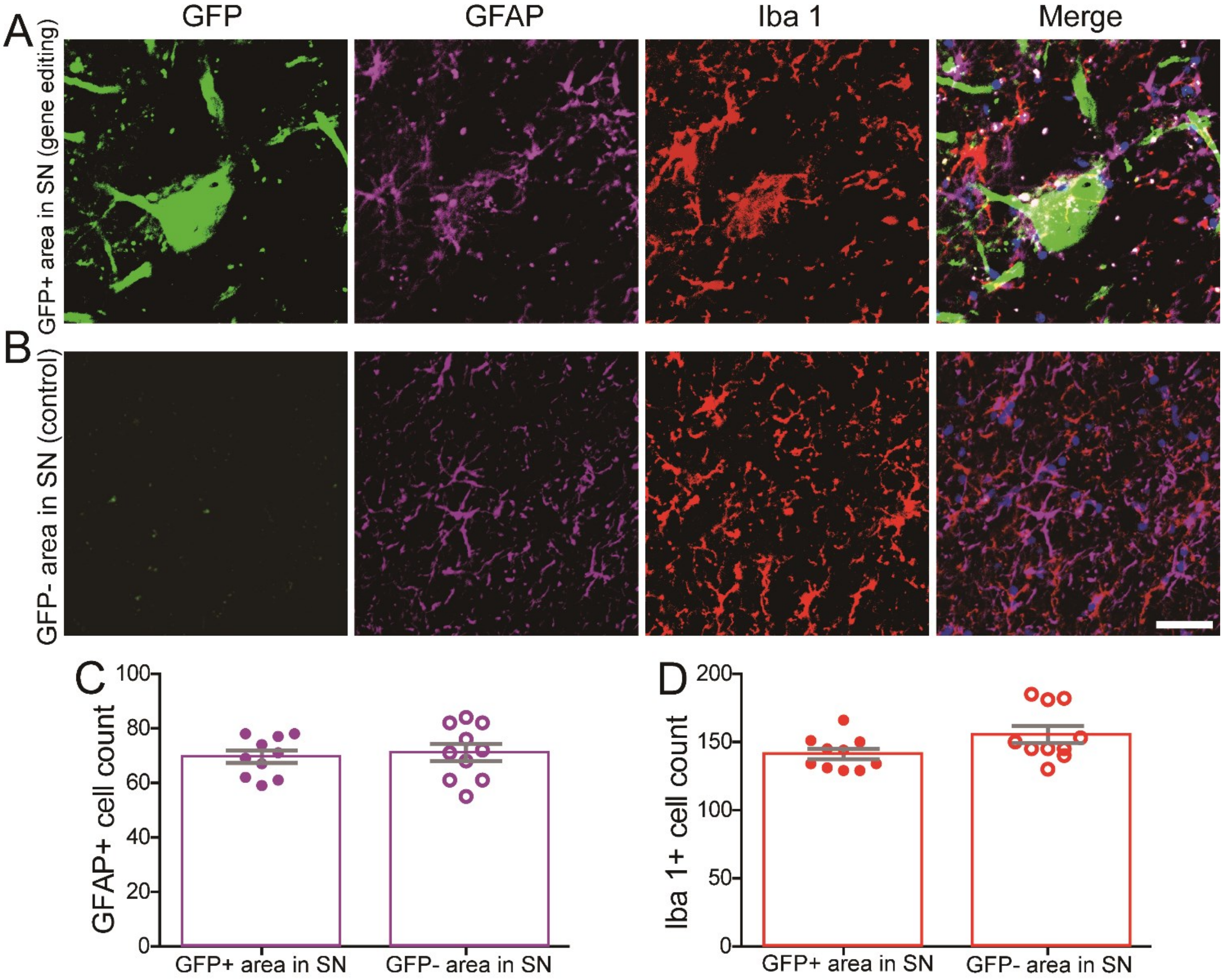


**Fig. S6. Glial cell immunostainings and quantifications.** (**A**) Astrocyte ($GFAP^+$ cell) and microglia ($Iba1^+$ cell) staining combined with GFP positive signals in gene-edited SN area. (**B**) Astrocyte and microglia staining combined with GFP negative signal in SN area (the control). (**C**) $GFAP^+$ cell count was not significantly increased (*P=0.687*). (**D**) Iba $1^+$ cell count in gene editing area was not significantly different from the control (*P=0.066*). Scale bar: 40 μm. Data are means ± SEM.

**Table S1.** Detail information for each monkey involved in this study.

| **Identification** | **Age#** | **Gender** | **Gene editing strategy** |
|---|---|---|---|
| 070229 | 10 | male | Left SN: *PINK1* & *DJ-1* |
| 070209 | 10 | male | Right SN: *PINK1* & *DJ-1* |
| 94089 | 23 | male | Left SN: *PINK1* & *DJ-1* |
| 97093 | 21 | male | Right SN: *PINK1* & *DJ-1* |
| 90067 | 26 | male | Left SN: *PINK1* |
| 95301 | 21 | male | Left SN: *PINK1* |
| 95075 | 21 | male | Left SN: *DJ-1* |
| 97081 | 19 | male | Left SN: *DJ-1* |

### indicates the age at death of each monkey.

**Table S2.** MRI-corrected coordinates for viral injections into SN of each monkey.

| | coordinates of bilateral SNs | |
|---|---|---|
| | **AAV delivered CRISPR/Cas9: *PINK1* & *DJ-1* editing side** | **AAV delivered CRISPR/Cas9: control side** |
| Monkey Young 1:**070229** | -4.80 (-6.80~-2.80) | -4.80 (-6.80~-2.80) |
| | +27.20 | -24.60 |
| | -36.70 | -34.50 |
| | 37˚ | 35˚ |
| Monkey Young 2: **070209** | -4.40 (-6.40~-2.40) | -4.40 (-6.40~-2.40) |
| | -26.00 | +25.70 |
| | -35.00 | -36.00 |
| | 41˚ | 38˚ |
| Monkey Old 1: **94089** | -4.35 (-6.35~-2.35) | -4.35 (-6.35~-2.35) |
| | +24.70 | -24.80 |
| | -36.80 | -37.00 |
| | 37˚ | 37˚ |
| Monkey Old 2: **97093** | 0 (-2.00~2.00) | 0 (-2.00~2.00) |
| | -23.90 | +25.20 |
| | -38.10 | -38.00 |
| | 35˚ | 37˚ |

**Coordinates description:** the optimum section of the coronal SN-distance before (-) or after (+) the marker plane (mm),

s: Left (+) or right (-) to the midsagittal plane (mm),

d: Distance (-) from the skull to the SN bottom (mm),

θ: Angle from the injection path to the vertical line (°).

**Table S3.** The improved Kurlan scale (a monkey-parkinsonism rating scale). Part A and part D are listed.

**Part A. Parkinsonian features (20 scores in total)**

**1.Tremor (L/R):** like the tremor of PD patients
- **0**-absent
- **1**-slight-low amplitude and only intermittently present
- **2**-moderate-moderate amplitude and present most of the time
- **3**-severe-high amplitude, virtually continuous, interferes with function

**2.Posture:** different from the postural instability of PD patients, not applicable to PD patients
- **0**-normal, erect
- **1**-stooped
- **2**-face down

**3.Gait:** partly like the postural instability of PD patients
- **0**-normal, use all four limbs smoothly
- **1**-walks slowly
- **2**-marked impaired, able to ambulate but very slowly and with effort
- **3**-severe decrease in ability to ambulate
- **4**-unable to ambulate

**4.Bradykinesia (generalized):** like the bradykinesia of PD patients
- **0**-normal speed and facility of movement
- **1**-mild slowing of overall movements
- **2**-moderate slowing of movements
- **3**-severe slowing of movements: slow, labored, and difficult to initiate and maintain movement
- **4**-unable to ambulate

**5.Balance:** partly like the postural instability of PD patients
- **0**-normal balance
- **1**-mild loss of balance on arising or with movement, holds onto cage for support
- **2**-major lapse in balance

**6.Gross motor skills (upper limb, L/R):** partly like the postural instability of PD patients
- **0**-normal, uses limb through a wide range of motion and activities
- **1**-noticeable decrease in capacity to use limb, but used consistently
- **2**-severe decrease in capacity to use limb, rarely used
- **3**-unable or refuses to use limb (including walking)

**7.Defense reaction (defensive and/or aggressive response to examiner):** not applicable to PD patients
- **0**-normal, reacts appropriately
- **1**-detectable impaired, slowed, abnormal or shortened response
- **2**-little or no response upon good provocation

**Part D. Clinical staging (5 stages)**
- **I.** Hemiparkinsonism, uses affected upper limb for walking
- **II.** Hemiparkinsonism, does not use affected upper limb for walking
- **III.** Bilateral parkinsonism, maintains adequate nutrition without assistance
- **IV.** Bilateral parkinsonism, requires assistance to maintain adequate nutrition, maintains upright position
- **V.** Bilateral parkinsonism, requires assistance to maintain adequate nutrition, face-down position

**Note:**

Part A was used to determine the Parkinsonian score while part D provided clinical measures of the Parkinsonian stage (Smith et al., Developing a stable bilateral model of parkinsonism in rhesus monkeys, 1993, *Neurosci*, **52**(1): p. 7-16).
